# Rapid evolution of coordinated and collective movement in response to artificial selection

**DOI:** 10.1101/2020.01.30.926311

**Authors:** Alexander Kotrschal, Alexander Szorkovszky, James Herbert-Read, Natasha I. Bloch, Maksym Romenskyy, Séverine Denise Buechel, Ada Fontrodona Eslava, Laura Sánchez Alòs, Hongli Zeng, Audrey Le Foll, Ganaël Braux, Kristiaan Pelckmans, Judith E. Mank, David Sumpter, Niclas Kolm

## Abstract

Collective motion occurs when individuals use social interaction rules to respond to the movements and positions of their neighbors. How readily these social decisions are shaped by selection remains unknown. Through artificial selection on fish (guppies, *Poecilia reticulata*) for increased social coordination (group polarization), we demonstrate that social interaction rules can evolve remarkably fast. Within just three generations, groups of polarization selected females showed a 15% increase in polarization, coupled with increased cohesiveness, compared to fish from control lines. They did not differ in physical swimming ability or exploratory behavior. However, polarization selected fish adopted faster speeds, particularly in social contexts, and showed stronger alignment and attraction responses to multiple neighbors. Our results demonstrate that animals’ social interactions can rapidly evolve under strong selection, and reveal which social interaction rules change when collective behavior evolves.

## Introduction

Moving animal groups display spectacular forms of coordinated behavior, with individuals moving together with high degrees of spatial and directional organization. This organization is often achieved by individuals using interaction ‘rules’ to respond to their neighbors’ movements and positions. For example, attraction, repulsion and alignment responses can act to maintain the cohesiveness and directional organization of groups (*1, 2*). The details of these interactions, and the social information individuals use to inform these decisions are now well described across many species (*3–7*). However, despite our growing knowledge of the mechanistic nature of social interactions in moving animal groups, we still know very little about the evolution of these social rules (*8, 9*).

For instance, while it has been established that intraspecific variation exists in animals’ social attraction and alignment towards conspecifics (*6, 10, 11*), it remains unclear whether such variation can be attributed to heritable differences in individuals’ social behavior, or is instead being driven by differences in individuals’ state, age, experience, or size (*12*). Indeed, while there are inherited differences in the tendencies of marine or benthic sticklebacks’ to school (*13*), those differences appear to be driven by genes affecting how social information is detected by neighbors (genes affecting the lateral lines system), and not necessarily how that information is behaviorally acted upon. Nevertheless, evolutionary models suggest that heritable differences in social decision-making should exist and persist in populations (*14*), and particular environments should favor particular social interactions depending on the selective forces present (*15*). What kinds of interactions are subject to selection, however, remains unclear.

In order to determine how selection can shape the social interaction rules that animals use to coordinate their movements, we performed a four-year artificial selection experiment using the guppy (*Poecilia reticulata*). Guppies are a model species for social behavior and evolution (*16*), and although they naturally shoal (*17*), their schooling tendencies tend to be weaker than in other species of fish, offering the potential for selection to increase social coordination. Our selection procedure targeted group polarization, a standard measure of directional coordination in animal groups. This metric captures the tendency of group members to align with each other’s directional headings. By artificially selecting for polarization over multiple generations we tested whether, and how quickly, coordinated group movement evolved when under strong directional selection. Importantly, polarization can only be measured in a group context but we nevertheless could apply an individual-level selection approach; our recently developed sorting protocol of repeated mixing and polarization-determination concentrates the individuals with the highest polarization propensities in few groups (*18, 19*). Those individuals could then be bred for the selection lines. Our artificial selection approach further allowed us to measure how selection shaped the social rules responsible for increased polarization in these groups.

Based on previous simulation and empirical studies, we had a number of *a priori* candidate mechanisms for how increased polarization could be achieved. These mechanisms include increased strength of alignment or attraction responses (*20, 21*), increased interaction ranges (*22*), increased number of influential neighbors (*23*), more frequent directional updating (*24*), faster speeds in social contexts (*5, 25*), or changes to individuals’ exploration or boldness (*10*). Here we identify which of these changes occurred to individuals’ social interaction rules following selection.

## Results

### The Artificial Selection Procedure

Across three independent selection lines (i.e. n = 3 replicate lines), we used a previously validated sorting method (*18, 19*) to identify the top 20% of female fish that consistently formed more polarized groups, and subsequently bred from those individuals. We focused on female behavior in our selection experiment because females of this species have a higher propensity to shoal than males (*17*). The sorting method involved open-field assays on groups of eight female fish (n = 16 groups per replicate), where fish were filmed when they explored an empty circular arena (diameter 550 mm, water depth 3 cm) together for ten minutes. Fish were then tracked using IDTracker (*26*) from the second to tenth minute, inclusive, from which the fish’ trajectories were subsequently analyzed. Across all frames in an assay, we calculated a group’s polarization (given by the total length of the sum of the eight unit vectors characterizing the orientation of each fish, divided by eight). Polarization scores closer to one indicate fish are oriented in the same direction, while scores closer to zero indicate fish are less aligned. After being assayed, the 16 groups were ranked for their median polarization scores, and half of each group’s members were subsequently swapped between adjacently ranked groups. This ranking and mixing of groups was repeated for 12 rounds, allowing us to create repeatable variation in polarization between groups (*19*). Twenty-six females from the four top-ranked groups in each line were then paired with unsorted males to breed the next generation of polarization-selected fish. To establish control lines (n = 3), we took 26 randomly selected females from the remaining groups, and bred from those fish. Once the progeny from each line were fully mature, the sorting method was performed again on the next two generations of females, providing a total of three generations of selection. Polarization and control line females were always paired with males from their own cohort. To ensure the control lines experienced the same experimental conditions as the polarization lines, control fish were placed in arenas and mixed between groups in the same way as the polarization lines, but were not sorted. For further details of the selection procedure see Fig. 1 and (*18, 19*).

**Fig. 1.**
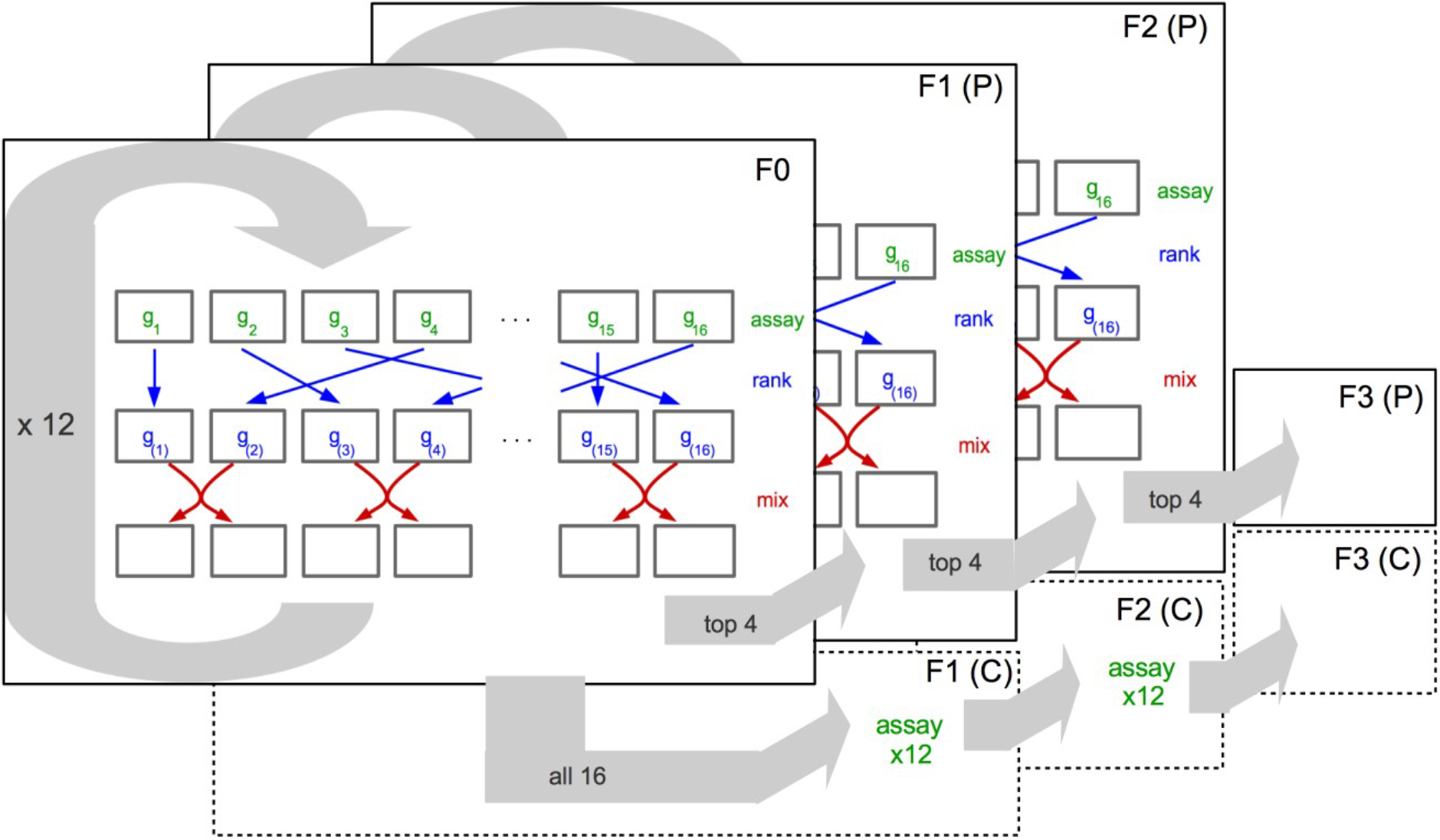
Schematic of one of three independent replicates of the selection experiment. Each layer represents a generation of females. Arrows within a layer illustrate the sorting procedure where we identified fish that formed the most polarized groups. To do this, groups were first assayed for their group polarization. Here variables *g*_1_ to *g*_16_ denote the 16 groups’ polarization scores in round *t*. These scores were subsequently ranked (blue arrows) with *g*_(1)_ to *g*_(16)_ denoting the ranked scores from lowest to highest. Following this, half of the group members were mixed with adjacently ranked groups (red arrows). This ranking and sorting procedure was repeated 12 times (circular grey arrow) before 26 fish from the top four ranked groups were bred for the polarization lines, and 26 fish from remaining 16 groups were bred for the control lines. This sorting procedure was repeated three times for the polarization lines (indicated by the layers), whereas fish from the control group experienced the same assaying and sorting, except fish from these control lines were not ranked.

### Evidence for Selection

We performed shoaling assays (as above) on the offspring of the polarization and control lines from generation three. We found that the polarization of groups across the three replicates was on average 15% higher in polarization lines (n = 88 groups) compared to control lines (n = 85 groups; difference replicate 1: 8.4%, replicate 2: 19.7%, replicate 3: 18.7%; LMM for all replicates: t = 6.45, df = 170, *P* < 0.001; Fig. 2A). Males did not display a significant response to selection (t=1.13, df=109, P=0.26), but weak differences between polarization and control lines existed in other behavioral measures consistent with the females. See Fig. S1 for results over all generations and discussion of the males.

**Fig. 2.**
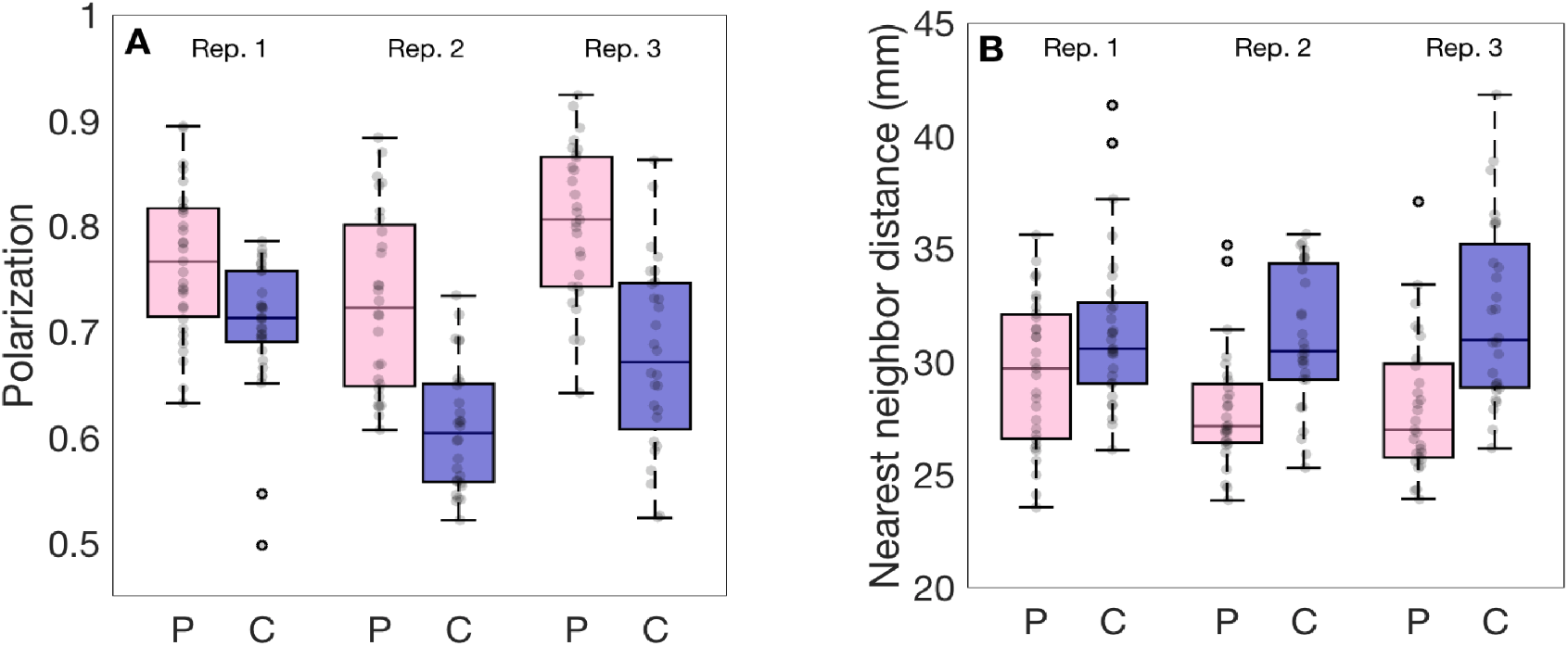
Polarization and nearest neighbor distance in groups of guppies artificially selected for polarization. Boxplots of (**A**) median polarization and (**B**) median nearest neighbor distance for groups of eight females in polarization selected (pink boxed) or control lines (blue boxes). Replicate lines 1, 2 and 3 are denoted above the boxes. Grey markers show individual data points (i.e. trials). Horizontal lines indicate medians, boxes indicate the interquartile range, and whiskers indicate all points within 1.5 times the interquartile range.

### Changes to Individuals’ Movement and Behaviour

We next tested whether selection had changed the movement characteristics of the fish in the polarization compared to control lines. As in many other fish species, guppies move with intermittent burst and glide phases (*15*), allowing us to characterize their movements in discrete steps (Fig. 3A, 3B). Groups of females from the polarization lines exhibited a 13.5 mm s^−1^ (26%) higher median speed in comparison to control lines (LMM: t = 5.59, df = 170, *P* < 0.001). We also performed open-arena assays on single fish and found that the difference in median speed between polarization and control lines was still significant, but less pronounced compared to the social context. Single fish from polarization lines were on average, 8.2 mm s^−1^ (17%) faster than single fish from control lines (LMM: t = 2.04, df = 117, *P* = 0.043). As speed is highly correlated with group polarization in shoaling fish (*10*), the increase in polarization seen in the selection lines could have been due to non-social selection for faster moving fish, or reduced swimming abilities in fish from the control lines. However, the polarization-selected lines were still 5.7% more polarized when controlling for median speed differences between the lines (LMM: t = 2.52, df = 169, *P* = 0.013); and there were no differences between the swimming abilities of fish in the polarization and control lines when tested for maximal swimming speed and endurance in a swim tunnel (LMM: t = −0.56, df = 64, *P* = 0.579; Fig. S2). Differences in behavior might also reflect differences between the polarization and control lines in overall ‘boldness’ or tendency to explore the arena, however, there were no differences in emergence time (i.e. ‘boldness’; LMM: t = −0.12, df = 28, *P* = 0.909; Fig. S3) or exploration (LMM: t = −0.38, df = 28, *P* = 704; Fig. S3) between the polarization and control lines when tested using a standard assay.

**Fig. 3.**
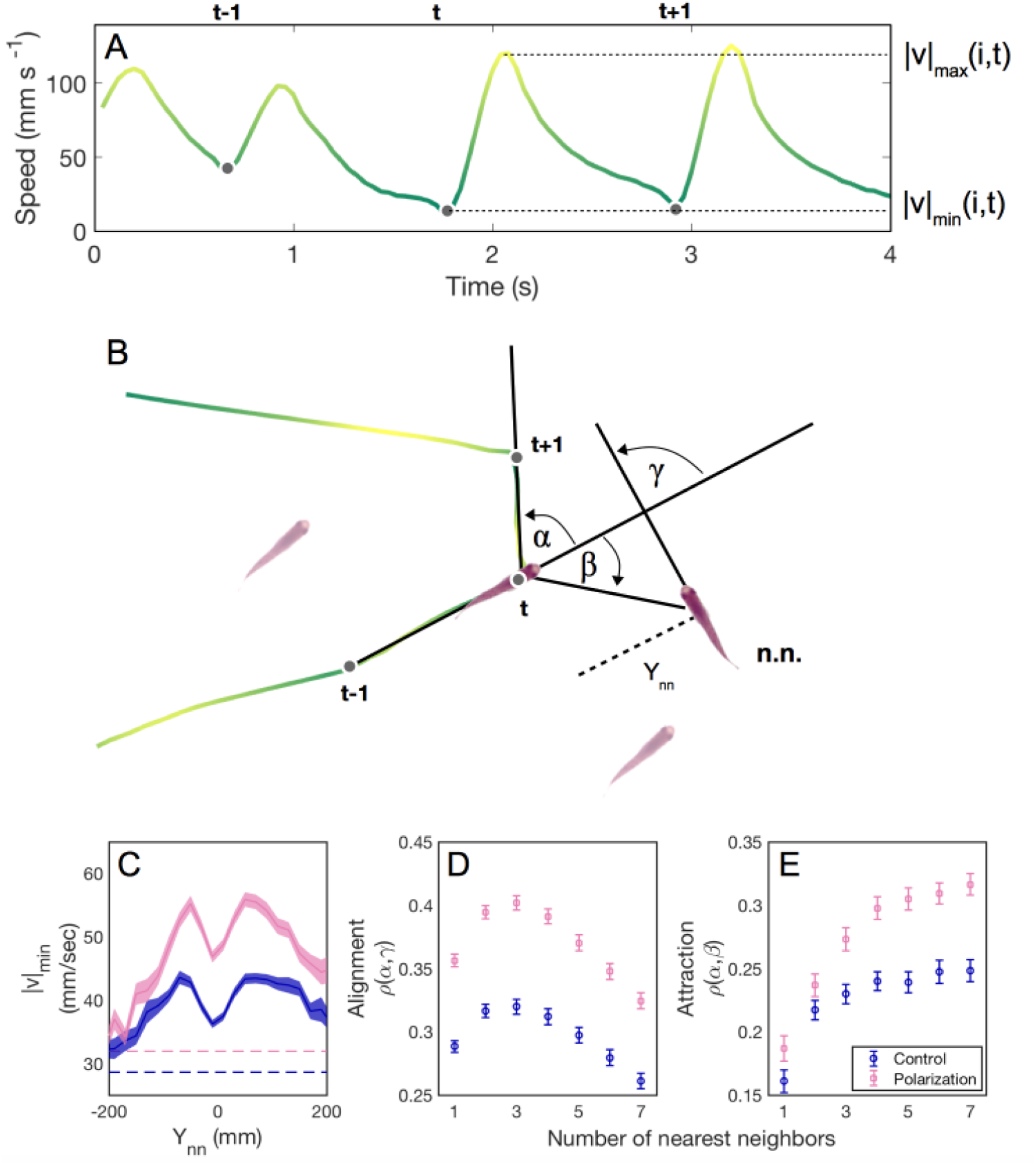
Burst and glide analysis and inferring social interactions of guppies artificially selected for polarization. (a) Time series containing three consecutive speed minima (dots) followed by bursting events. (b) The corresponding trajectory for fish *i* (in the center). The positions at the preceding and following speed minima are used to calculate the turning angle α of fish *i* at time *t*. In this example, the turn has the same sign as the nearest neighbor orientation (i.e. alignment) γ, and the opposite sign to the attraction angle β. (c-e) The social interactions of females in the control lines (blue) and polarization selection lines (pink) in response to nearest neighbors. (c) The mean speed minimum when the nearest neighbor is in front (+) or behind (−) by a distance *Y*_*nn*_. The error region shows the standard error in the mean over trials. (d) Alignment and (e) attraction responses to the geometric center of *k* nearest neighbors, where *k* ranges from 1 (nearest neighbor) to 7 (all conspecifics). The Spearman correlations ρ were computed for each *k* and for each trial for all β and γ with absolute values of less than 90 degrees. The set of β was additionally restricted to time points where the *k* neighbors were all less than 200 mm from the focal fish (*Supplementary Materials*). Means (symbols) and standard errors (bars) were calculated for each selection line from these correlation coefficients.

In order to further investigate whether the social environment affects the speed that fish adopted in the polarization and control lines, we identified the speeds at which fish decided to accelerate (|v|_min_; Fig. 3A) and plotted this as a function of the distance to their nearest neighbor (Fig. 3C). We found that while fish from the polarization lines generally maintained higher speeds than fish from control lines, these differences were particularly apparent when fish were close to their neighbors, with differences in speed between the lines becoming less pronounced as neighbors moved further apart. This provides further support that differences in speed were, at least in part, modulated by interactions with conspecifics. Polarization and speed results were also robust when controlling for potential differences in thigmotaxis (‘wall-hugging’, i.e. propensity of swimming close to the walls) between the lines (*Supplementary Materials*).

### Selection on Individuals’ Social Interaction Rules

Polarization lines were significantly more cohesive than control lines (Fig. 2B; 3 mm, or 10% smaller median nearest neighbor distance; LMM: t = −5.5, df = 170, *P* < 0.001), a finding that would not be expected if there were changes to individuals’ speeds but not in their social interactions (*10*). We therefore tested whether selection had altered the social interaction rules of the polarization compared to control lines. Many models and subsequent empirical work have identified that fish, including guppies, use attraction and alignment responses to coordinate their movements (*5, 15*). To test whether selection had changed the strength of these alignment or attraction rules, we first extracted the turning angles α (Fig. 3B) that a fish made between its movement bursts. We then calculated the Spearman rank correlation over an entire trial between turning angles (α) and nearest neighbor directions (β) to quantify attraction strength, and with neighbor orientations (γ) to quantify alignment strength (*Supplementary Materials*). The strength of these correlations, therefore, acts as a proxy for the strength of these interactions. We found that fish from polarization lines had on average 23% higher correlations between turning angle and nearest neighbor orientation, and hence stronger alignment responses (LMM: t = 9.91, df = 170, *P* = 0.007; Fig. 3D). There was also a non-significant trend for polarization lines to have stronger attraction towards their nearest neighbor than control lines (LMM: t = 1.94, df = 170, *P* = 0.054). When we included speed as a covariate in our models, fish from the polarization lines still showed 9% higher alignment strength than fish from control lines (LMM: t = 4.82, df = 169, *P* = 0.016; see also the *Supplementary Materials*), showing that increased alignment responses were not only due to faster motion.

We then asked whether selection had changed the number of neighbors individuals were responding to during these attraction and alignment responses, using the centroid and mean orientation of the *k* nearest neighbors to calculate β and γ, respectively (*Supplementary Materials*). The shape of the alignment and attraction responses, measured as a function of the number of influential neighbors, was qualitatively similar for both lines, declining after three to four neighbors in the case of alignment and plateauing at three to four neighbors in the case of attraction (Fig. 3D, 3E). This finding is reminiscent of the rule structuring in zonal models of collective motion, where alignment interactions occur with closer neighbors, and attraction responses with more distant neighbours (*27*). Although polarization-selected fish were not significantly more attracted to their nearest neighbor than control fish, attraction strength to multiple neighbors was stronger in the polarization than control lines. Attraction strength to k = 7 nearest neighbors (i.e. the group centroid) increased in the polarization lines by 27% compared to control lines (LMM: t = 4.56, df = 170, *P* < 0.001; Fig. 3E; see *Supplementary Materials* for male results). Selection may also have acted on the distance over which these alignment or attraction responses occurred. This is a common parameter in metric-based models of collective motion (*21, 27*), and we tested it by analysing occasions when the nearest neighbor was in front of a focal individual, and the focal individual either turned towards that neighbor with an attraction response (|α−β| < 30 degrees) or turned to align with that neighbor with an alignment response (|α−γ| < 30 degrees). We took the distance at which these responses occurred more frequently than by chance as a proxy for their interaction range (*Supplementary Materials*). We found no conclusive evidence that there were solid differences in either the attraction (LMM: t = 1.95, df = 170, *P* = 0.053) or alignment ranges (LMM: t = −1.6, df = 170, *P* = 0.11) between the selection and control lines.

## Discussion

Our results confirm that social interactions in a collective motion context are heritable, and that they can be rapidly shaped by directional artificial selection, leading to more polarized and cohesive groups. In particular, our selection regime changed three important aspects of individual behavior: 1) speed, 2) the strength of the alignment response, and 3) the attraction strength to larger groups of conspecifics. Below we discuss the implications of these discoveries for our understanding of the interaction rules that lie behind evolutionary changes in collective motion.

First, increased speed has been suggested as an important and relatively simple mechanism behind more coordinated collective motion behavior (*5, 15, 25*). Importantly in our assays, the observed speed differences between the polarization and control lines were strongest in social contexts. For instance, the speed differences between lines were most prominent when close to conspecifics, suggesting that social facilitation may play an important role in how speed affects the increased polarization (*28*). Moreover, the observed differences in alignment were robust (albeit smaller) also when controlling for speed in the analysis, and we did not find any differences between polarization and control lines in our assays of physical swimming ability and behavioral stress responses. Hence, although our results indicate that speed changes play an important role for the behavioral differences between the polarization and control lines, we propose that such changes require a social context to be important for evolutionary shifts in collective motion.

After controlling for speed differences between the lines, fish from polarization lines were still more likely to align with their neighbors’ directional heading than fish from control lines. The number of neighbors, or the range over which these alignment responses occurred, however, was not different between the selection lines. These results are consistent with how social responsiveness is often implemented in theoretical models of collective motion, where individuals weigh the tendency to travel in their own goal-orientated directions against the adoption of neighbors’ directional headings (*1, 29*). It is possible, therefore, that selection acted on intrinsic differences in the social responsiveness of individuals, as has been predicted to exist in wild populations (*14, 30*). As well as increased alignment responses, polarization lines also showed stronger attraction responses to multiple conspecifics. While increased attraction is typically viewed in context of reducing predation risk through selfish herd effects (*15, 31*), increased attraction to others can also be viewed in the context of social decision-making, where individuals are often attracted towards larger numbers of neighbors (*32*). Our results suggest, therefore, that selection may have acted on how individuals weigh social information, ultimately leading to differences in group structure and social dynamics.

One possible explanation of the substantial change in polarization in only three generations is that the traits under selection have a simpler genetic background than is usually assumed. Examples do exist where seemingly complex behavior, such as burrowing behavior in mice, can have a relatively simple genetic architecture (*33*). In our experiment, however, the selection regime has changed several aspects of interaction rules in the polarization lines. We therefore view a very simple genetic background to these differences as unlikely, unless that architecture has pleiotropic effects across all these rules. Another explanation is due to the behaviors under selection here being a product of interactions between the behavior of a focal individual and the other individuals in the group. Such social interactions have been suggested to be strongly influenced by indirect genetic effects, where expression of a trait in one individual alters expression of the trait across the social group, thereby amplifying the effect of selection (*34*). Indirect genetic effects could have played a role also in our experiment and increased the response to selection, but more work is needed to reveal the genetic architecture behind the observed differences.

In nature, which social rules evolve will ultimately depend on the selective forces present. Previous research has suggested that selective forces including the social environment (*14*), resource availability or distribution (*8, 35*), and predation risk (*9, 15, 22*) are likely to shape individuals’ alignment and/or attraction responses. In turn, the social responses that evolve will have functional consequences for groups’ abilities to track environmental gradients (*36*) and transfer information about detected threats or resources between group members (*37, 38*). However, to fully understand the evolution of these social rules, we also need to better understand the costs associated with evolving them. We found that increased coordination and cohesive behavior was associated with increased energy expenditure (i.e. increased speed). Similar energetic costs of coordination have been reported in flocks of birds (*39*). Future analyses on the polarization selection lines will investigate the costs and benefits of increased coordinated and collective movement in ecologically relevant settings.

It is noteworthy that the response to selection on polarization was weaker in males than in females. We specifically selected on female collective behavior in our experiment, and this could explain the weaker response in males. However, behaviors with strong fitness effects should have strong inter-sexual genetic correlations. The profound ecological differences between males and females in the guppy, with females having much higher propensity of shoaling (*40*), could explain the sex differences we observe. Our results certainly suggest that the genetic correlation between males and females for polarization behavior is relatively low, possibly due to differences in genetic architecture for social behavior between males and females (*41*).

In summary, our research has identified the social interaction rules that are affected by directional selection on polarization, and shown that such traits are susceptible to fast evolutionary changes. An integrated approach to understanding social behavior through artificial selection combined with detailed behavioral measurements now offers considerable opportunities to understand the evolution and maintenance of social decision-making and collective behavior.

## Material and Methods

### Collective motion analysis

Collective motion analyses were performed on 2565 mins of videos, obtained via a Point Grey Grasshopper 3 camera (FLIR Systems, resolution 2048 × 2048 px, frame rate 25 Hz), in MATLAB R2017b (details on video processing and data extraction can be found in (*19*)). Speed minima and maxima were found by first smoothing the speed profiles of individual fish (using a Savitzky-Golay filter degree three, span 12) and applying the *findpeaks* function. We found that turns in the trajectories typically came three frames (0.12 s) after a speed minimum, and accordingly applied this delay when calculating turning angles. For the assays of eight and single fish exploring the arenas, one of each measure was extracted per nine minute trial. The median distance from the edge of the arena was log-transformed and the polarization was transformed using an inverse logistic function.

### Swimming speed and boldness tests

To test if selection changed the physical swimming ability of the fish, we measured the critical swimming speed of fish in a flow chamber (*42*) in 66 females and 62 males from the control and polarization selection lines. The flow chamber consisted of a 115 cm long transparent PVC pipe with an inside diameter of 1.8 cm through which aerated water was pumped at controllable speed (‘swim tunnel’). We measured critical swimming speed by subjecting fish to increased velocity tests: the guppies were forced to swim against a current which was increased in discrete steps, until exhaustion occurred and they were swept against the outflow end of the tube. After a 2-minute acclimation period at a low velocity of 6.5 cm s−1, we increased the current velocity by 2.2 cm s−1 every 30 s until the guppy reached exhaustion and was unable to detach itself from the outflow mesh for 3 seconds. Temperature was maintained at 25.0°C ±1.5°C. Results are shown in Fig. S2 and Table S1.

We also measured boldness and exploration of 60 females and 60 males in a standard emergence test; in a 50 l tank with 3 cm of water. The starting compartment (20 × 10 cm) was separated from the exploration compartment (20 × 40 cm) by an opaque partition with an eight cm wide opening. After two minutes of acclimation in the starting compartment an opaque trap door was lifted to allow access to the exploration compartment. The time it took until individuals left the starting compartment was used as indicator of boldness and the number of 5 × 5 cm plots visited (15 in total, every time a new plot was visited, this was added to the total area visited) was used as indicator for exploratory tendencies 3. Non-emerged fish after the maximum time of 10 minutes were removed from the analysis (15 from each set of female lines, two from male polarization lines, four from male control lines). The time to exit a shelter was used as a measure of a fish’s boldness, and the area explored by each fish was used as a measure of their exploratory tendencies. Both measures were log-transformed. Results are shown in Fig. S3 and Table S1.

### Statistics

We tested for differences between selection lines using linear mixed-effect models. Separate models were used for individual trials and groups of eight, as well as for males and females. Selection line was incorporated as a fixed effect. For tracked motion assays, mean body size (estimated from IDTracker) was incorporated as a covariate, as well as median speed when controlling for activity. Replicate was used as a random effect for the intercept and the selection effect. Normality of residuals were checked using Kolmogorov-Smirnov tests, with a maximum KS statistic of 0.107. Residuals were plotted against fitted values to visually check for correlations and heteroscedasticity. Analyses were done in MATLAB R2017b.

## General

We thank Anna Rennie, Eduardo Trejo and Annika Boussard for help with fish husbandry.

## Author contributions

NK, KP, and DS conceived the idea to the study. AK, AS, MR, SDB, JHR, HZ, KP DS, and NK designed the details of the selection procedure. MR set up the filming apparatus, AK performed and AS analyzed the data of the selection experiment. AK, AF, LSA, ALF, and GB performed and AK, AS and SDB analyzed the data of the behavioural assays. AK, AS and JHR created the figures. AK, AS, JHR, JEM, NIB, DS, and NK wrote the manuscript. All authors contributed to the final version of the manuscript.

## Data availability

All data and code will be deposited in Dryad.

## Supplementary Information

### Artificial Selection

Fig. S1 shows the polarization of the original F0 female groups, and the selection and control lines in subsequent generations.

### Additional collective motion analysis

#### Complimentary burst and glide results

Female groups from the polarization lines made more frequent bursts than the corresponding control lines (mean time between bursts was shorter by 0.022 s or 2.7%; LMM: *t* = 3.01, *df* = 170, *P* = 0.003) and made smaller turns (0.042 radians or 6.9%; LMM: *t* = 5.79, *df* = 170, *P* < 0.001). This was most likely due to the negatively correlated biomechanics of speed and turning combined with the arena’s geometry.

#### Thigmotaxis analysis

To control for potential differences in thigmotaxis (attraction to the walls of the arena), that could affect the potential for polarization if one group tended to swim nearer the edges, we analyzed the median distances from the edge of the arena across the selection lines. The polarization line females were on average 2.6 mm further from the center of the arena in trials with single fish (LMM: *t* = 2.12, *df* = 117, *P* = 0.036) compared to the single fish controls (257.9 mm), and 3.1 mm further from the center of the arena in groups (LMM: *t* = 3.49, *df* = 170, *P* < 0.001) compared to the control groups (253.9 mm). We therefore repeated our analysis only including frames from the videos where the mean distance from the arena edge was more than 50 mm (i.e. less than 225 mm from the center). This analysis showed that polarization line females were still 10.2% more aligned than the control lines (LMM: *t* = 4.34, *df* = 170, *P* < 0.001), demonstrating that differences in thigmotaxis were not driving the differences in polarization between the polarization and control lines.

#### Conspecific speed

Attraction and alignment turning responses to the nearest neighbor (quantified by the Spearman rank correlation, see Fig. S4) varied with the neighbor’s speed and position. To illustrate this, we plotted aggregated attraction and alignment responses as a function of both forward distance to nearest neighbor and speed of this neighbor at the bursting time of the focal fish (Fig. S5, S6). Attraction responses were strongest when the neighbor was close in front, while alignment responses were strongest when the neighbor was close in front and travelling more quickly. By subtracting the control line heat maps from the polarization line heat maps (Fig. S5 C, F), it is clear that the female fish from the polarization lines show an increased response in these regions. That is, for comparable speeds and positions, the polarization line females show an increased correlation between turning response and the nearest neighbor’s position and orientation, lending further support that social interactions in the females differed between selection lines. This was less apparent in the males (Fig. S6 C, F), although these patterns occurred in the same direction as the females.

#### Multiple neighbor responses

We used both metric and topological models to test turning responses to multiple neighbors. The topological attraction model used the mean position of the nearest *k* neighbors as the X variable for the Spearman correlation. See Fig. S4 for a typical trial for *k* = 1 and *k* = 7. In all per-trial correlations, only data points with *X* < 90 degrees were used, as the dependence is monotonic in this region. The topological attraction correlations for females as a function of *k* are shown in Fig. S7 A.

In the metric attraction model, the X variable is the mean position of all neighbors within a distance *r*, rather than a specific number of neighbors. As can be seen in Fig. S7 B, this correlation reached a peak around 200 mm and then declined. This weakening of the response to distant neighbors is consistent with the decline in the topological attraction model for large *k*. Restricting the topological model to neighbors within 200 mm hence partially controls for differences in cohesion between lines. In this case, the topological attraction reached a plateau at *k* > 3 for both lines (see main text Fig. 3 D, E).

In the topological and metric alignment models (Fig. S7 C, D), the X variable is the mean orientation of the neighbors within the first *k* neighbors or radius *r* respectively. The orientation of the focal fish was not included in this calculation.

#### Range calculations

We calculated the range of a response (A) using kernel-smoothed distributions of nearest neighbor distances, *R*. A typical response range for A can be defined as a range of *r* satisfying

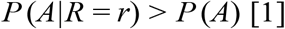

which by Bayes’ rule is equivalent to

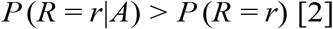

Hence, for each trial, we calculated these two probability density functions of the nearest neighbor distance, *R*. The density function corresponding to *P* (*R* = *r*) is calculated using all decision points in which the neighbor is in front of the focal fish. All nearest neighbor distances at these points are input to a kernel-smoothed density estimator with bandwidth of 10 mm. The density function *P* (*R* = *r A*) is calculated in the same way, but with the extra restriction that there is a response A (i.e. the turning angle is within 30 degrees of the vector corresponding to the neighbor position (attraction) or orientation (alignment)). We then modify the above expression slightly to define the range *r*_max_ as:

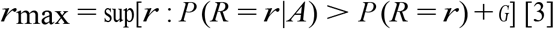

We used a small constant offset *G* = 2.5 × 10^−4^ to account for small fluctuations in the tails of the distributions (see Fig. S10).

### Male analysis

When testing offspring from the third generation of selection, we did not find any significant differences in the polarization of male groups between the polarization (*n* = 56) and control lines (*n* = 56), although trends were in the same direction as the females (LMM: *t* = 0.944, *df* = 109, *P* = 0.26). While males did not significantly differ in polarization between the lines, in concordance with the females’ results, males’ median speed was higher by 5.7mm s^−1^ (9%) in polarization compared to the control lines when tested in groups (LMM: *t* = 2.95, *df* = 109, *P* = 0.004), but not when tested alone (LMM: *t* = 0.708, *df* = 109, *P* = 0.48). The nearest neighbor alignment responses and the group attraction responses were respectively 10% and 22% stronger in males from the polarization lines compared to control lines (nearest neighbor alignment LMM: *t* = 2.76, *df* = 109, *P* = 0.007; group attraction LMM: *t* = 3.21, *df* = 109, *P* = 0.002; Fig. S9), again, a finding that was consistent with results from the females. Together, these results suggest that although males from polarization and control lines differed in speed and social interactions in the same way as female fish, this did not generate the same strong differences in polarization that were observed in the females. This may be due to the generally reduced social tendencies of males compared to females, which could be due to differences in social behavior that are sexually linked.

---

**Fig. S1.**
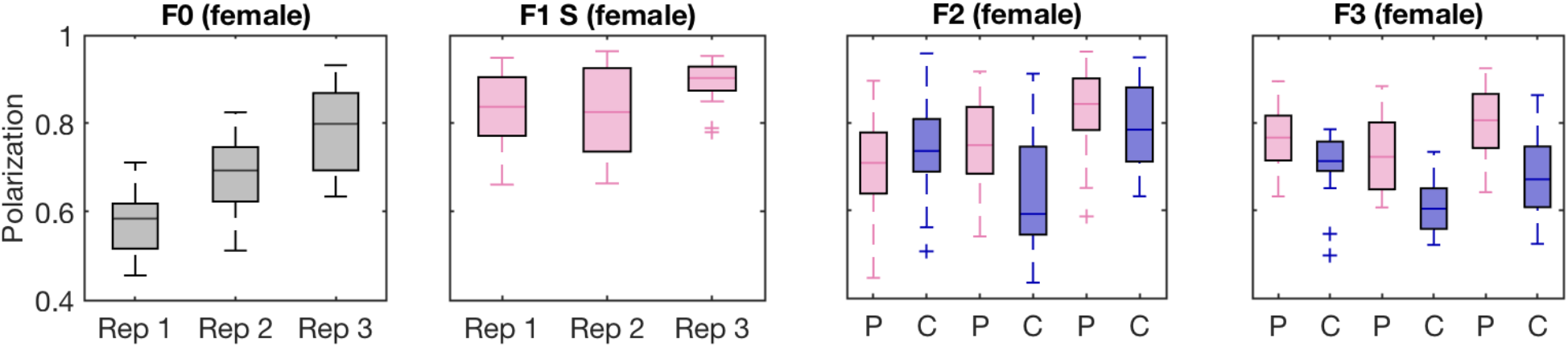
Polarization of females in F0 (base population), F1 (first sorting round, control lines not filmed), F2 and F3. Higher polarization values indicate more coordinated schooling behavior. Horizontal lines indicate medians, boxes indicate the interquartile range, and whiskers indicate all points within 1.5 times the interquartile range. P: Polarization line (pink bars), C: Control line (blue bars).

**Fig. S2.**
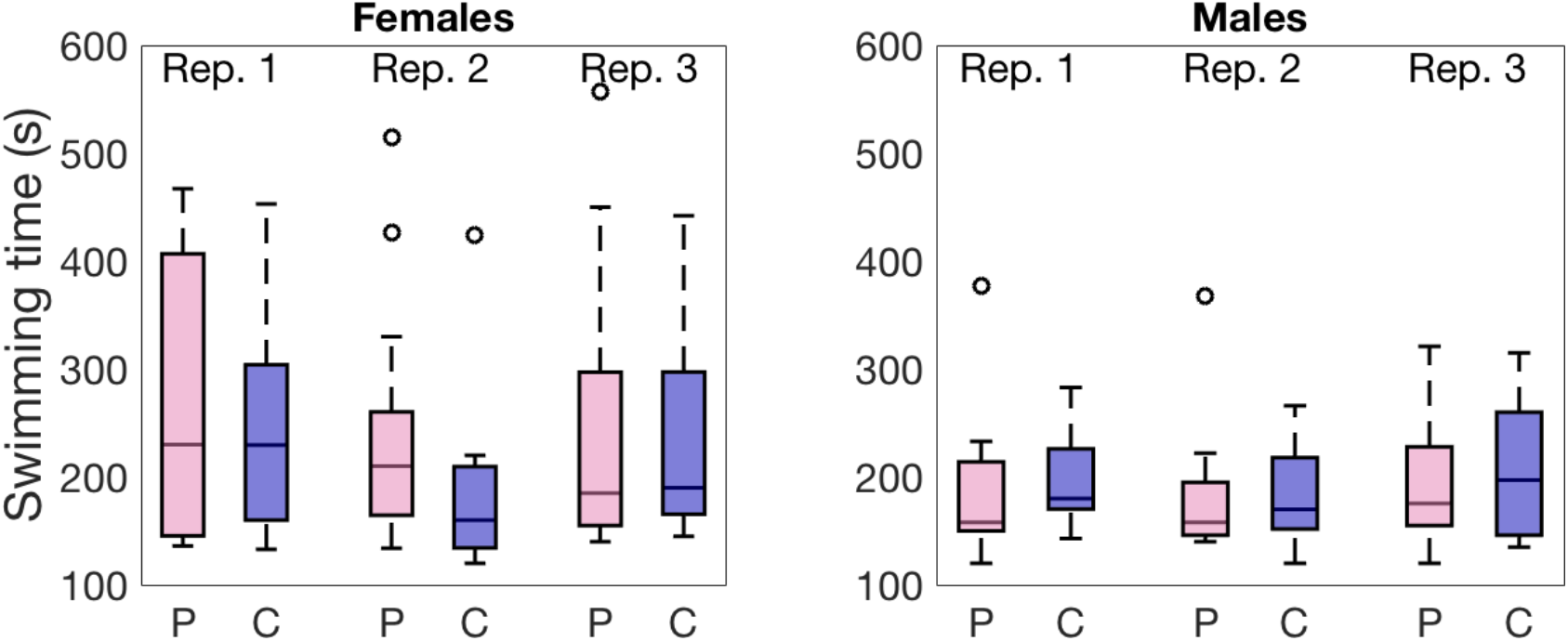
In a swim tunnel with increasing laminar water current polarization (P, pink) and control (C, blue) lines showed similar maximal swimming speeds in females (left panel) and males (right panel). See Table S1 for statistics on log-transformed variables. Note that swimming time and speed are equivalent as swimming speed was increased at constant time intervals. Horizontal lines indicate medians, boxes indicate the interquartile range, and whiskers indicate all points within 1.5 times the interquartile range.

**Fig. S3.**
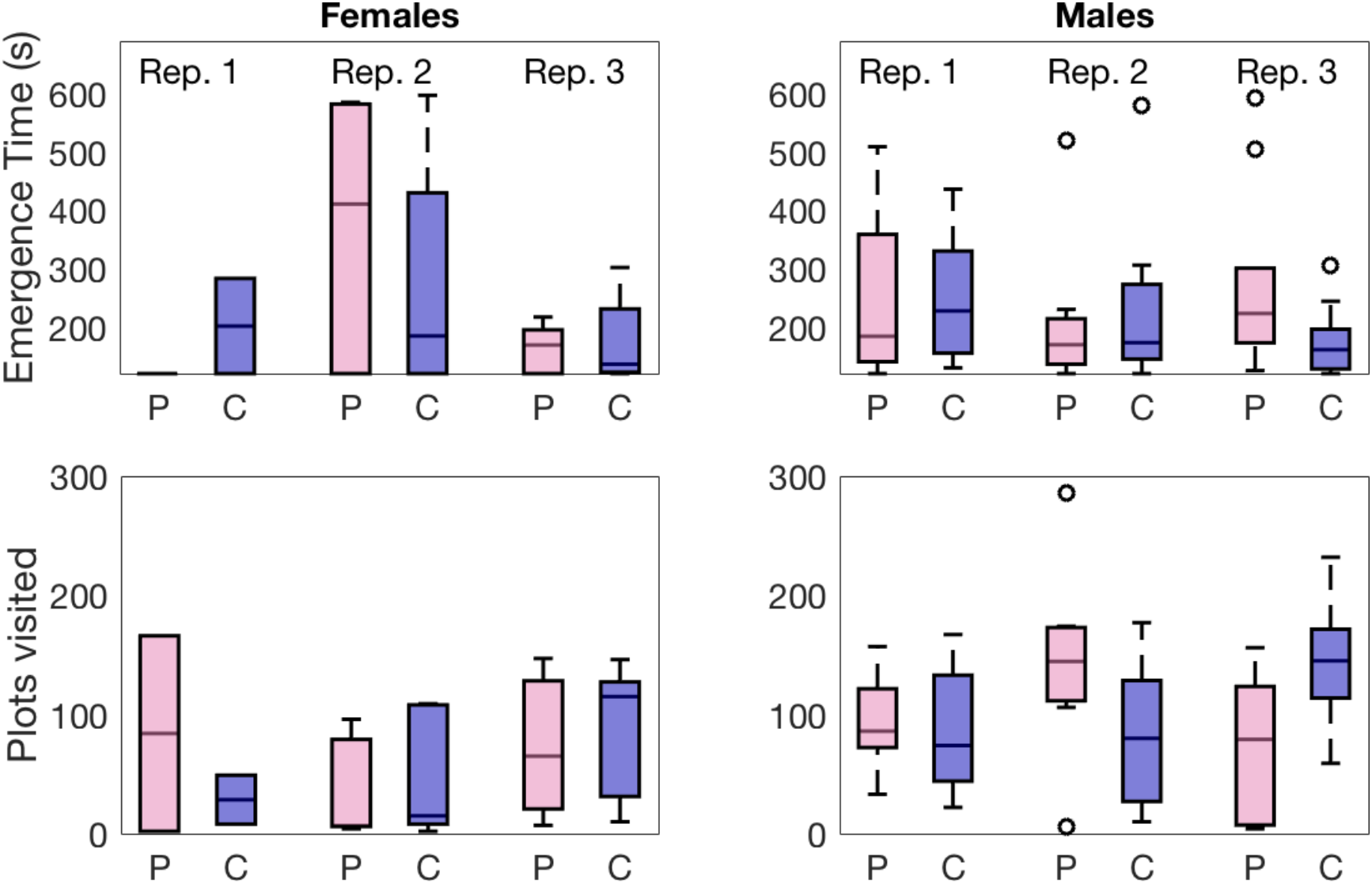
An emergence test revealed no differences in emergence time (‘boldness’ top panel) or the number of plots visited (‘exploration’ bottom panel) between polarization (P, pink) and control (C, blue) lines in females (left panels) or males (right panels). After log-transformation, no differences were found in females or males (see Table S1). Horizontal lines indicate medians, boxes indicate the interquartile range, and whiskers indicate all points within 1.5 times the interquartile range.

**Fig. S4.**
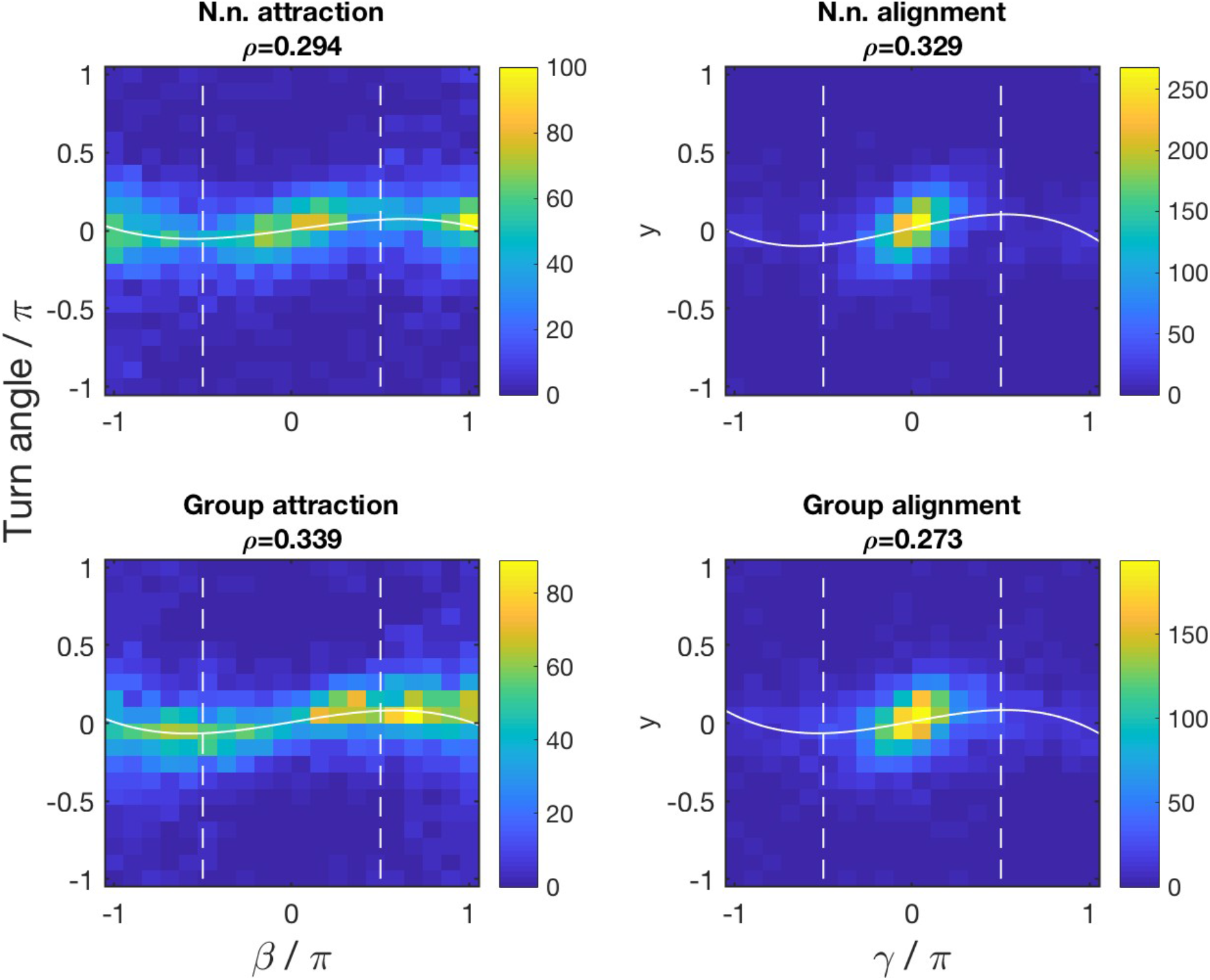
Rank correlations for a single female trial. Each panel is a bivariate histogram of the turning angle (vertical axis) against a predictor (horizontal axis). The color bar shows the number of data points in each bin. A third-order polynomial is fit to each set of points and is shown as a solid white line. The Spearman correlations shown above each panel are calculated from all points where the absolute predictor angle is less than 90 degrees (i.e. within the dotted white lines).

**Fig. S5.**
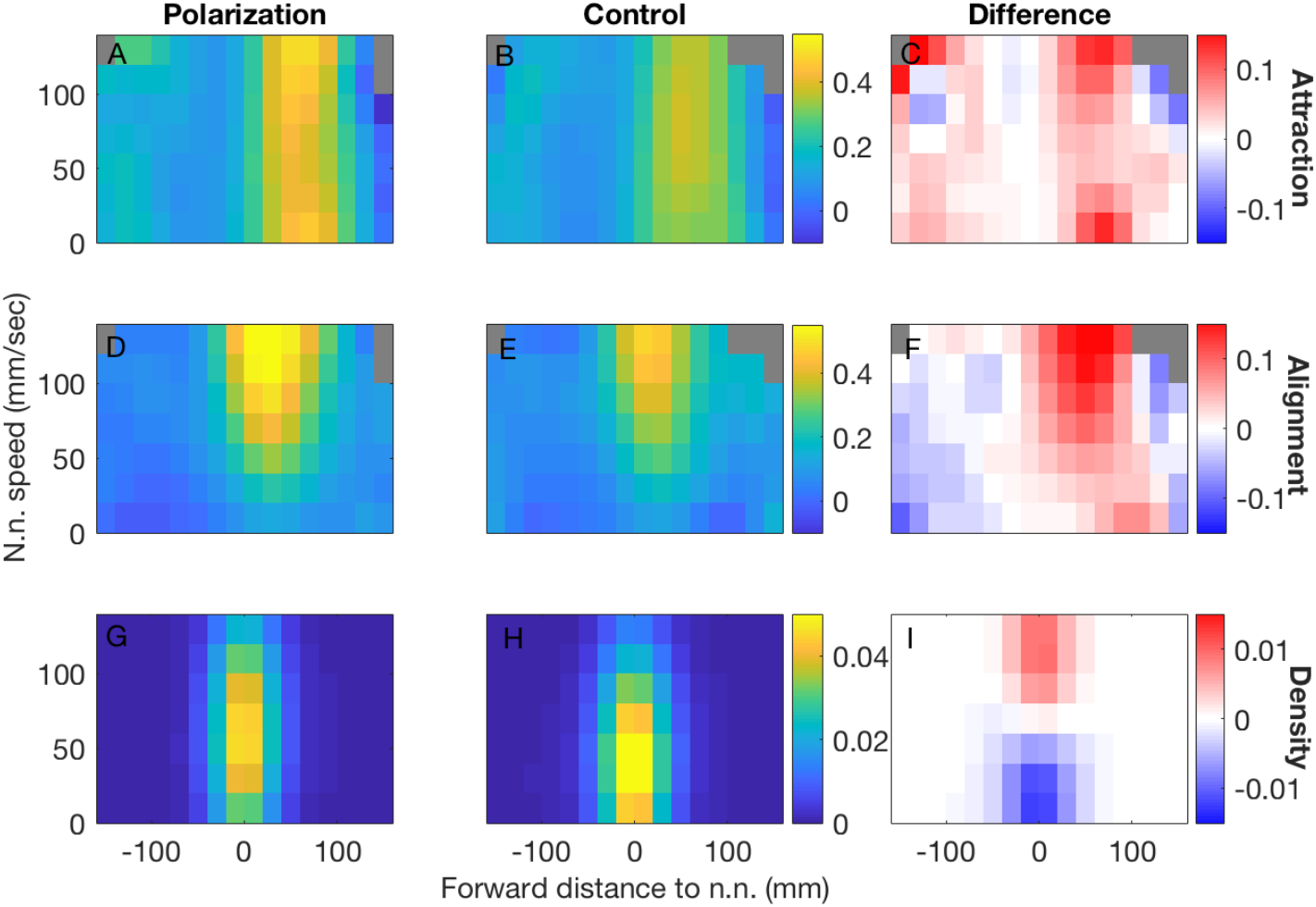
Typical burst and glide behavior of females as a function of nearest neighbor position and speed. In all panels the horizontal axis is the forward distance to the nearest neighbor and the vertical axis is the nearest neighbor speed. In each bin within panels A, B, D, and E, the Spearman rank correlation was calculated between the turning angles and the vector corresponding to the position or orientation of the nearest neighbor to quantify the attraction or alignment strengths respectively. Panels C&F show differences in rank correlation between the two panels to the left. Panels G&H show the density of data points, while panel I shows the difference in densities.

**Fig. S6.**
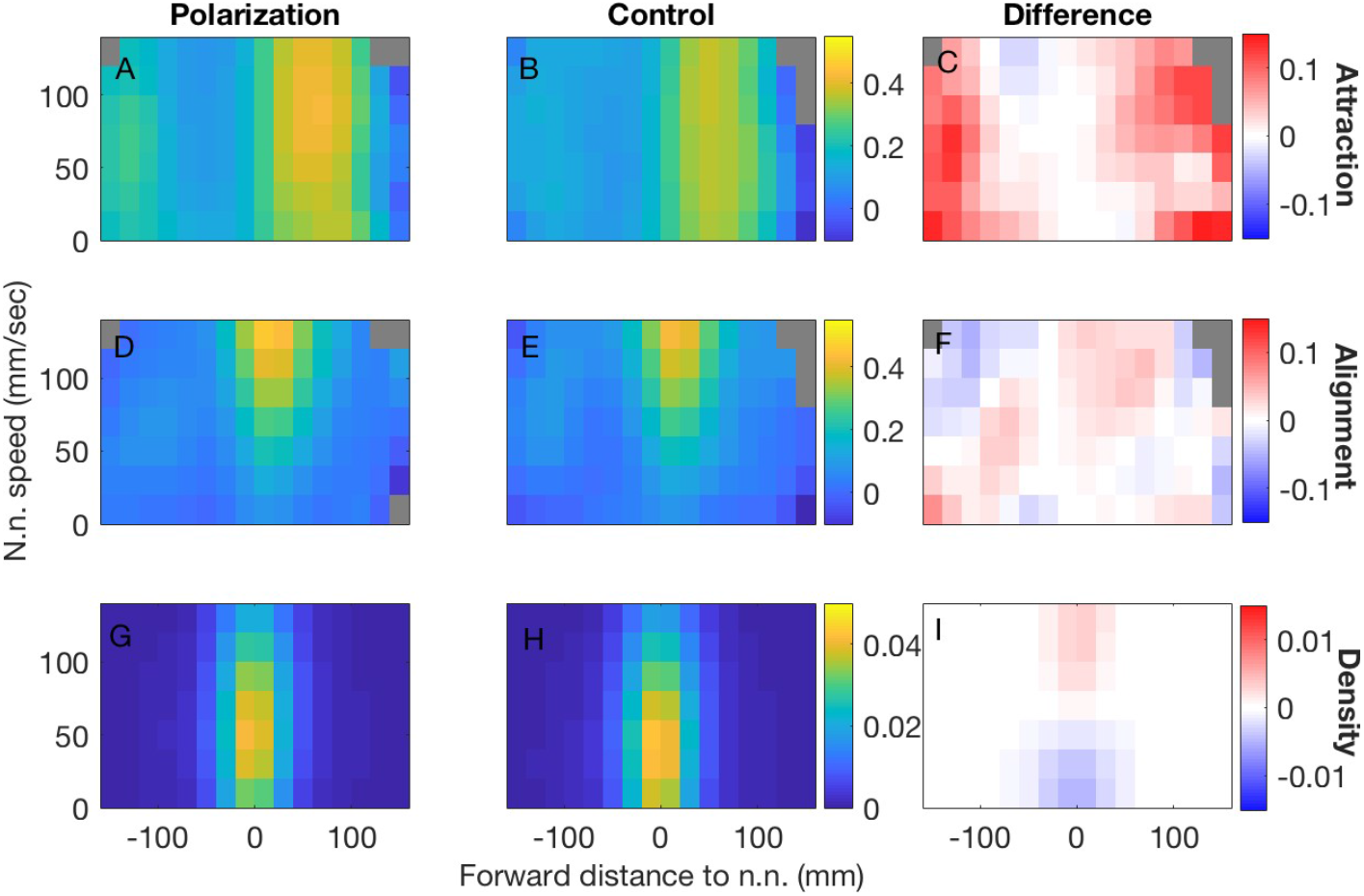
Typical burst and glide behavior of males as a function of nearest neighbor position and speed. All panels as in Fig S4. In all panels the horizontal axis is the forward distance to the nearest neighbor and the vertical axis is the nearest neighbor speed. In each bin within panels A, B, D, and E, the Spearman rank correlation was calculated between the turning angles and the vector corresponding to the position or orientation of the nearest neighbor to quantify the attraction or alignment strengths respectively. Panels C and F show differences in rank correlation between the two panels to the left. Panels G and H show the density of data points, while panel I shows the difference in densities.

**Fig. S7.**
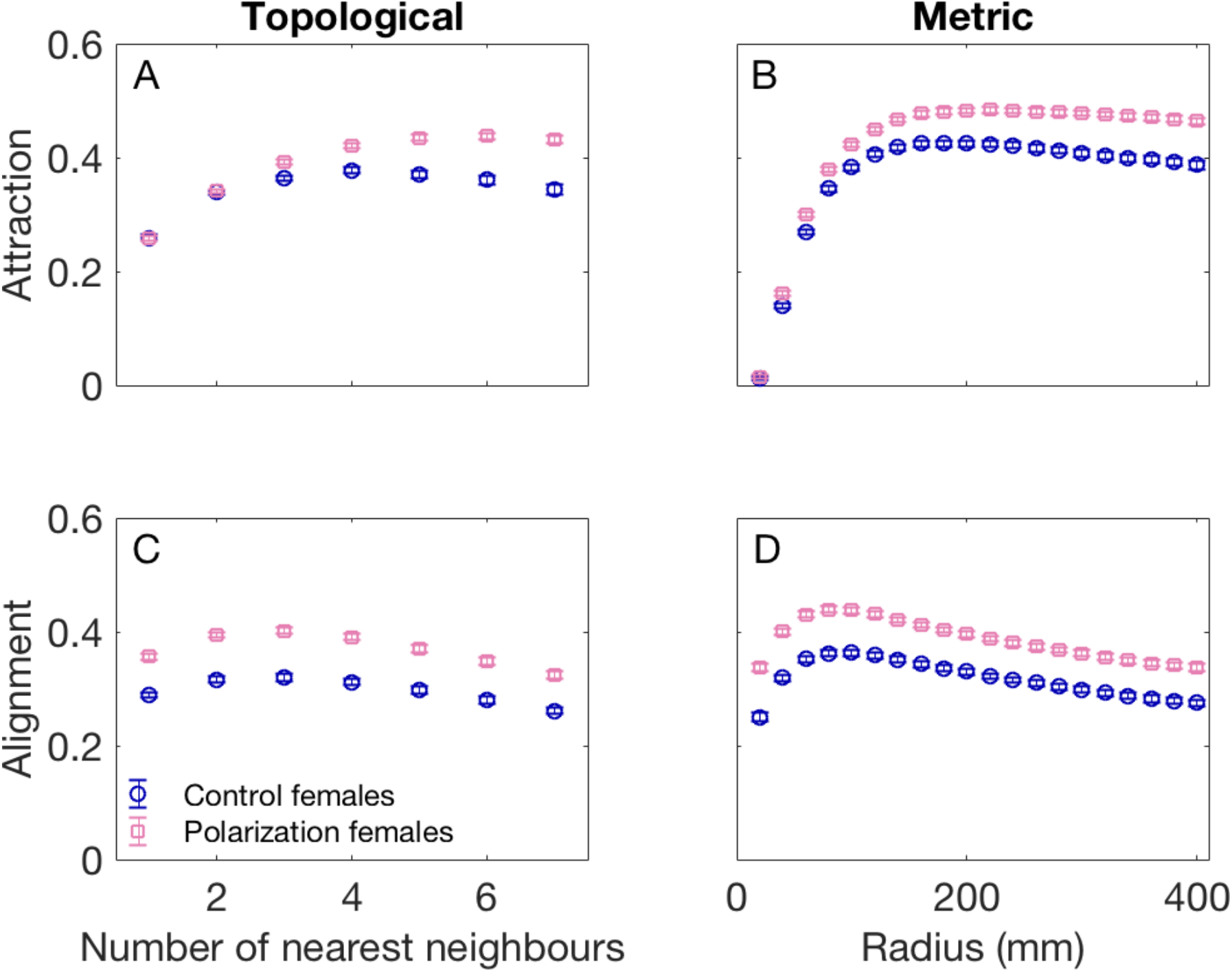
Female alignment and attraction responses vs. topological distance (A, C) and metric distance (B, D). Shown are means of the per-trial rank correlations for the selection and control lines; with error bars indicating the standard errors over trials.

**Fig. S8.**
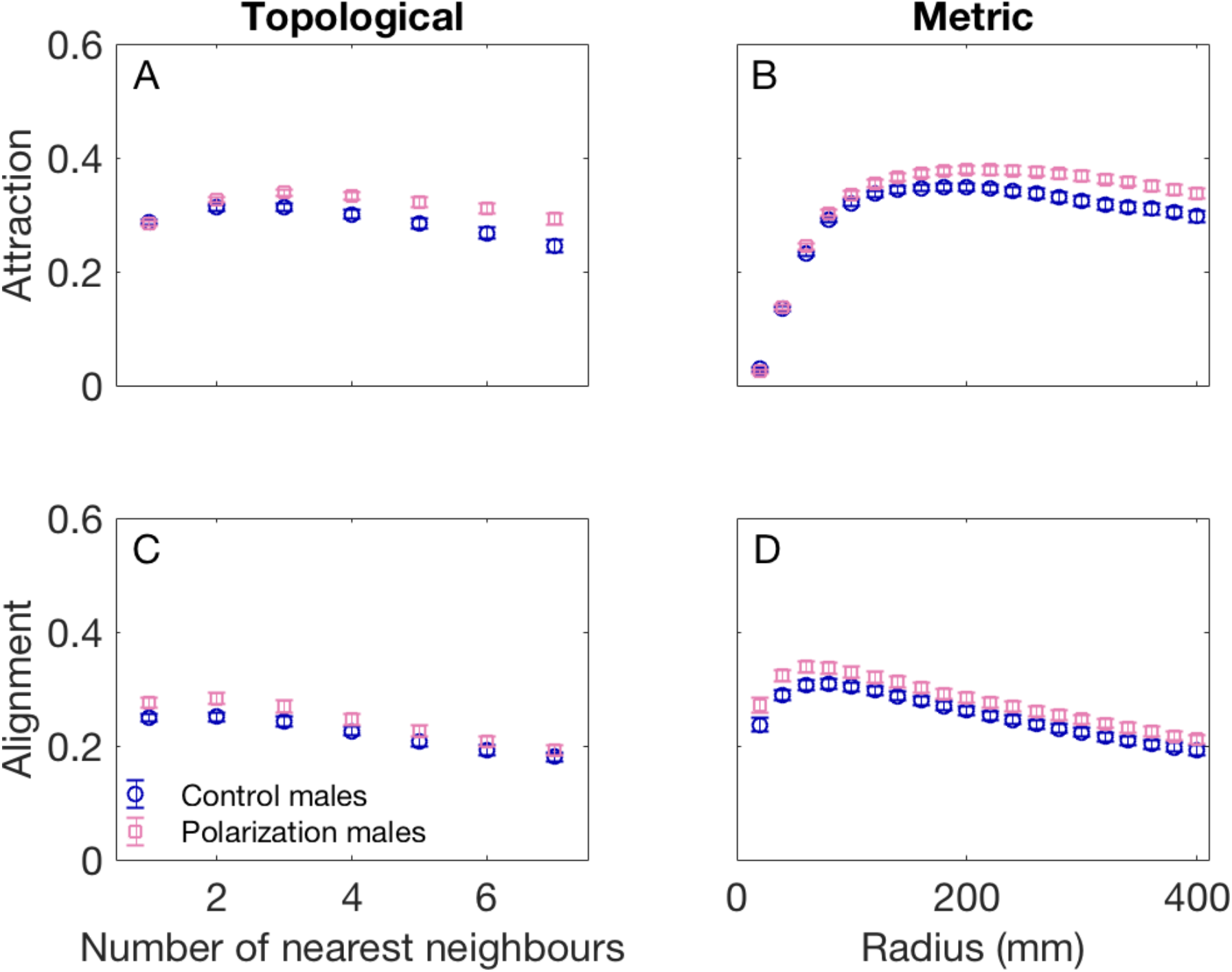
Male alignment and attraction responses vs. topological distance (A, C) and metric distance (B, D). Shown are means of the per-trial rank correlations for the selection and control lines; with error bars indicating the standard errors over trials.

**Fig. S9.**
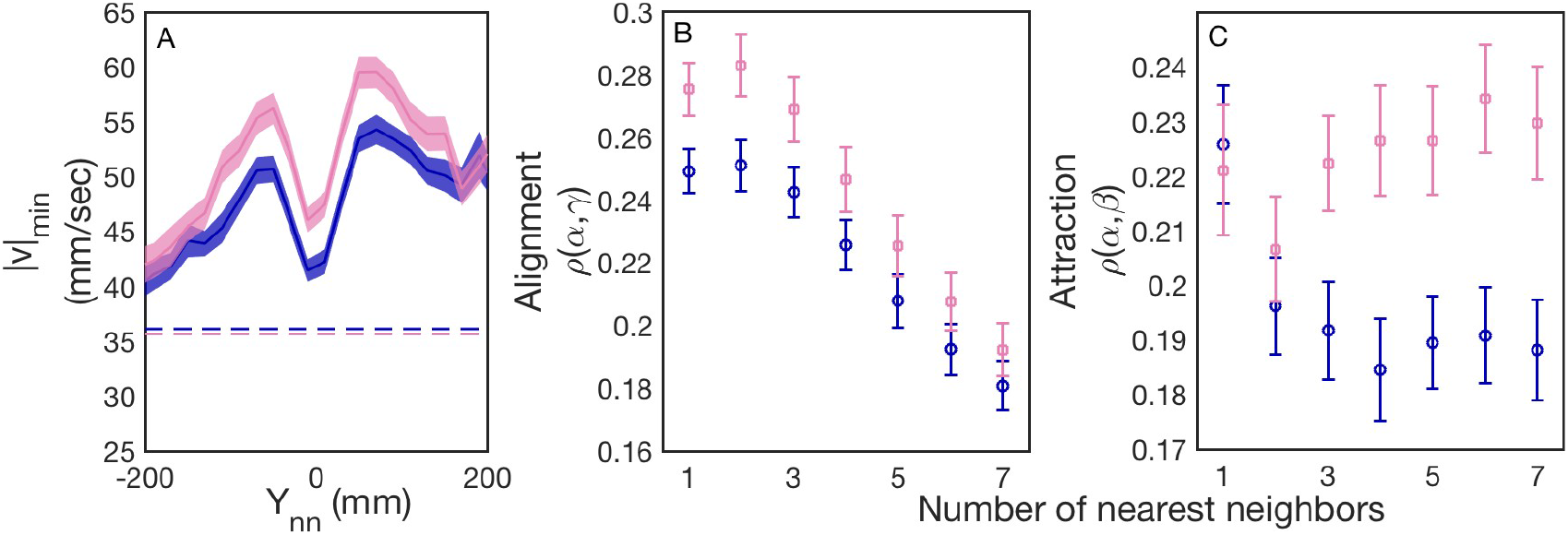
Typical burst and glide behavior of males in the control lines (blue) and polarization selection lines (pink) in response to nearest neighbors. (A) Mean speed minimum when the nearest neighbor is in front (+) or behind (−) by a distance *Y*_*nn*_. The error region shows the standard error in the mean over trials. (B) Alignment and (C) Attraction responses to the geometric center of *k* nearest neighbors, where *k* ranges from 1 (nearest neighbor) to 7 (all conspecifics). The Spearman correlations *ρ* were computed for each *k* and for each trial for all *β* and *γ* with absolute values of less than 90 degrees. The set of *β* was additionally restricted to time points where the *k* neighbors were all less than 200 mm from the focal fish. Means (symbols) and standard errors (bars) were calculated for each selection line from these correlation coefficients.

**Fig. S10.**
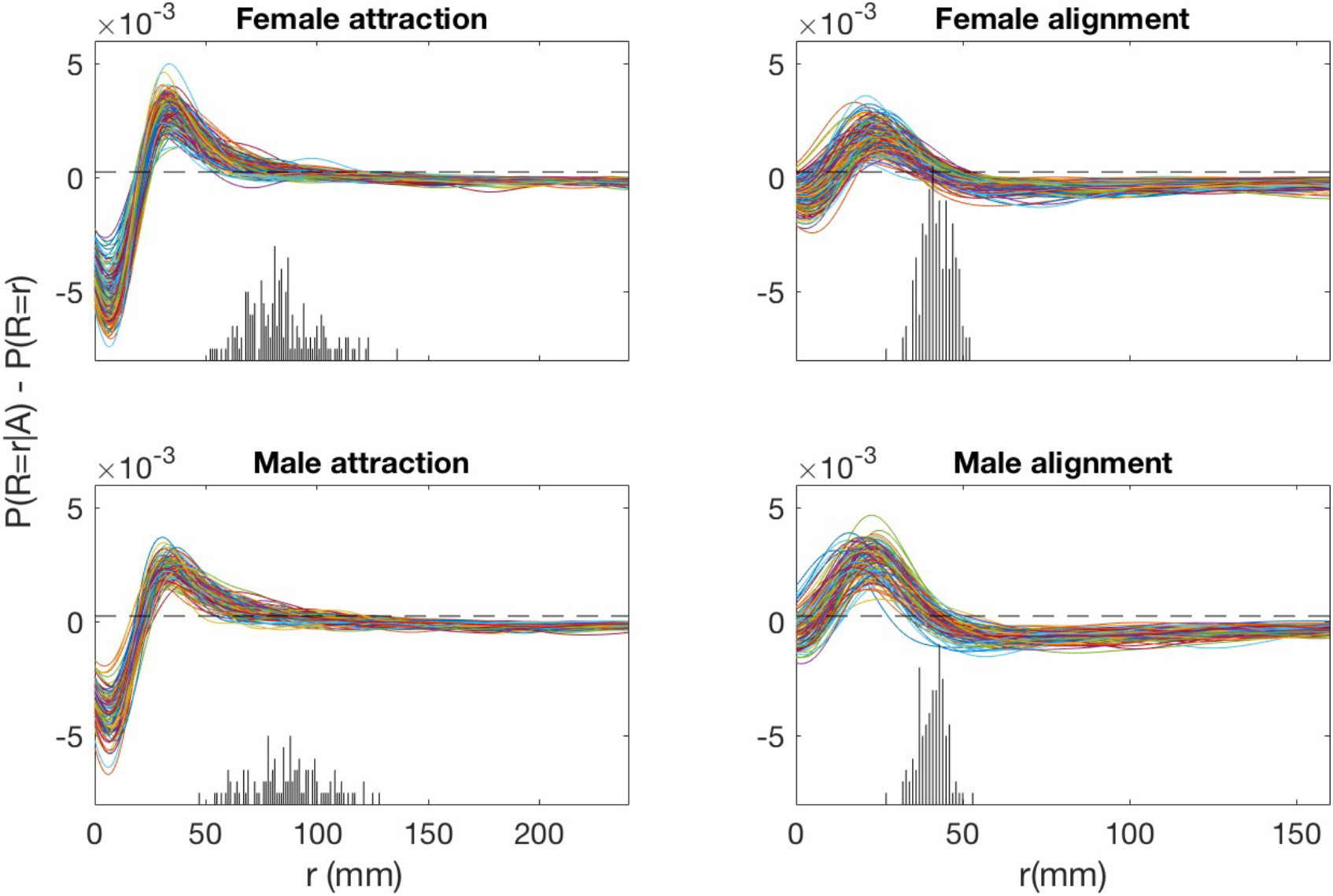
Calculation of attraction and alignment ranges. In each panel, each curve corresponds to the difference in kernel-smoothed distributions for one trial of eight fish. A region where the curve is above zero indicates a response above the average level. The dotted line shows the cut off at *s*. The histograms below show the distributions of the final points in *r* above the cutoff, which are used as the attraction and alignment ranges.

**Table S1.**
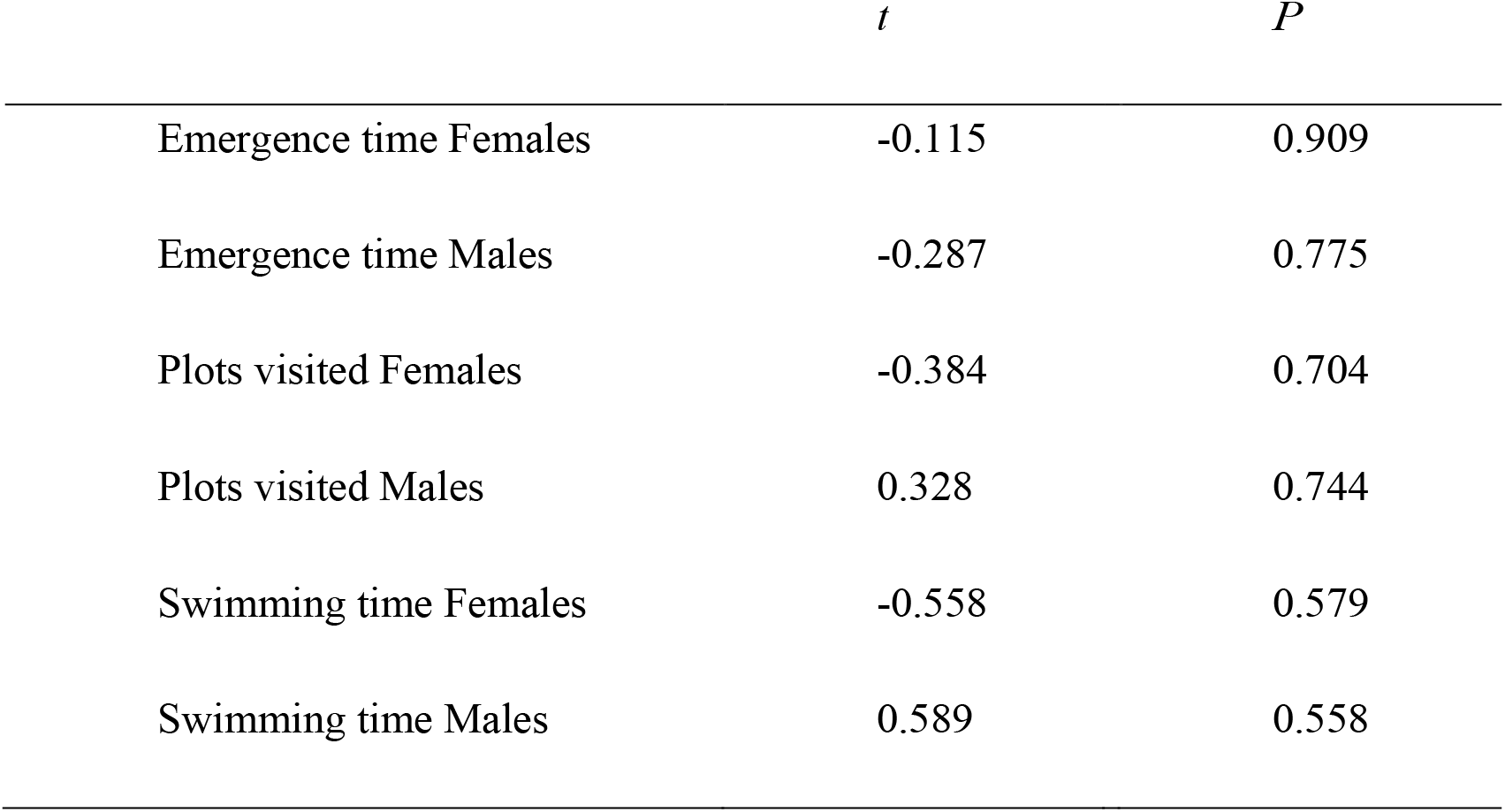
Results from the fitted linear mixed-effect models on emergence time, plots visited, and swimming time.

|  | *t* | *P* |
| --- | --- | --- |
| Emergence time Females | -0.115 | 0.909 |
| Emergence time Males | -0.287 | 0.775 |
| Plots visited Females | -0.384 | 0.704 |
| Plots visited Males | 0.328 | 0.744 |
| Swimming time Females | -0.558 | 0.579 |
| Swimming time Males | 0.589 | 0.558 |

